# Agent-based computational modeling of the stochastic dynamic behavior of actin filaments recapitulates the homeostatic cortical array in plant epidermal cells

**DOI:** 10.1101/2025.06.06.658236

**Authors:** June Hyung Kim, Weiwei Zhang, Anjali Iyer-Pascuzzi, Christopher J. Staiger, Taeyoon Kim

**Affiliations:** Weldon School of Biomedical Engineering, Purdue University, West Lafayette, IN 47907, USA; EMBRIO Institute, Purdue University, West Lafayette, IN 47907, USA; Department of Biological Sciences, Purdue University, West Lafayette, IN 47907, USA; Department of Botany and Plant Pathology, Purdue University, West Lafayette, IN 47907, USA; Faculty of Science and Technology, Keio University, Kohoku Ward, Yokohama 223-0061, Japan

## Abstract

The homeostatic cortical actin array in plant cells plays important roles in fundamental processes, including intracellular transport, secretion, cell expansion, and cytoplasmic streaming. In response to diverse chemical and mechanical signals, the cortical array can remodel within minutes to assume new configurations or altered filament abundance. The homeostatic cortical actin array of some plant epidermal cells comprise sparsely distributed individual actin filaments and actin bundles, which enables tracking and quantitative analysis of dynamic properties over many minutes at high spatiotemporal resolution. Previous studies using quantitative live-cell imaging, small molecule inhibitors, and genetic mutations reveal the robust dynamic steady state of the cortical actin array, with individual filaments showing a behavior termed stochastic dynamics. Compared to experimental findings, computational efforts focused on the plant actin cytoskeleton are lacking, although computational models have the potential to define underlying mechanisms of actin array homeostasis and remodeling. Here, we used an agent-based computational model to reproduce the stochastic dynamic behavior of individual actin filaments in epidermal cells with consideration of key governing factors, including the nucleation, polymerization, depolymerization, severing, capping, and branching of filaments. Our model was able to reproduce experimental observations with respect to the abundance and length of filaments as well as the rates or frequencies of dynamic events. This model can be used to study the role of myosin motors and other actin-binding proteins, as well as the effects of signaling events and fluxes in cellular second messengers, on actin dynamics in plant cells.

## INTRODUCTION

The actin cytoskeleton in plant cells plays important roles in cell morphogenesis and expansion, cytoplasmic streaming, defense response, vesicle trafficking, and intracellular reorganization (Szymanski & Staiger, 2018; Yuan et al., 2023). The plant actin cytoskeleton exhibits highly dynamic behaviors which are known to be coordinated by a myriad of accessory proteins (Staiger et al. 2009; Smertenko, Deeks, and Hussey 2010; Henty-Ridilla, Li, et al. 2013; Blanchoin et al. 2010). Interaction between thermally fluctuating actin monomers in the ATP state (i.e., ATP-G-actin) rarely results in the formation of a stable nucleus consisting of three monomers, a process known as de novo nucleation (Blanchoin et al., 2014; Goode et al., 2023; Pollard & Borisy, 2003). Individual actin filaments (F-actin) adopt an asymmetric structure with assembly occurring predominantly at the barbed end with ATP-G-actin. After assembly, actin subunits in the filament undergo ATP hydrolysis followed by Pi release, which subsequently generates ADP-F-actin that is prone to severing and disassembly from the pointed end (Kudryashov & Reisler, 2013). The organization and turnover of actin arrays are determined by the cooperation of actin with a variety of actin-binding proteins, some of which enhance the actin turnover cycle. Actin Depolymerizing Factor (ADF)/cofilin promotes severing and consequently increases the depolymerization rate by binding to ADP-F-actin (Henty et al., 2011; McGough et al. 1997; Maciver and Hussey 2002; McCullagh, Saunders, and Voth 2014). Heterodimeric capping protein binds to filament barbed ends, thereby preventing polymerization and filament-filament annealing (S. Huang et al., 2003; J. Li et al., 2012). Additionally, experimental findings indicate that a majority of severed ends are capped shortly after severing, suggesting an interplay between severing and capping proteins (Staiger et al., 2009). On the other hand, adenylyl cyclase-associated protein increases the rate of nucleotide exchange on G-actin, providing a sufficient supply of ATP-actin for the assembly of actin filaments (Chaudhry et al. 2007).

Profilin is an actin-binding protein that preferentially binds to ATP-G-actin (Krishnan and Moens 2009). Its association with actin occludes a crucial interaction site required for de novo nucleation (Blanchoin et al., 2014; Goode et al., 2023; Pollard & Borisy, 2003). Due to ∼2–3 fold higher profilin concentration relative to total actin concentration in the plant cytoplasm, the majority of the actin monomers are predicted to be bound to profilin (Chaudhry et al. 2007; Pollard, Blanchoin, and Mullins 2000). The limited pool of profilin-free monomers substantially reduces the frequency of de novo nucleation, which likely contributes to the sparse actin array seen in the plant cortex (Sun et al. 2018; Paul and Pollard 2008; Kovar, Drøbak, and Staiger 2000; Kovar et al. 2006).

Slow de novo filament nucleation is compensated by nucleation factors (Pollard, 2007). The Arp2/3 complex binds to the side of actin filaments to create a pseudo nucleus with two actin-related protein monomers at a characteristic angle of ∼70° (Amann & Pollard, 2001; Goley & Welch, 2006; Mehidi et al., 2021; Mullins et al., 1998). In *Arabidopsis*, the inhibition of Arp2/3 complex shows abnormal morphology of epidermal cells with obvious defects in cytoskeletal organization and function, such as cell wall assembly and auxin distribution (S. Li et al., 2003; Pratap Sahi et al., 2018; C. Zhang et al., 2013). Although the importance of Arp2/3 complex as a nucleator is clear, the exact mechanism by which it coordinates actin array organization and turnover in plants remains elusive. Formins are another class of nucleation factors that consist of two formin-homology (FH) domains. In general, the family of formins show a wide variety of behaviors with varying elongation and nucleation rates (Kovar et al. 2006; Moseley and Goode 2005; Michelot et al. 2006). The FH2 domain processively associates with the barbed ends of actin filaments and promotes enhanced polymerization by recruiting profilin-bound actin, resulting in the formation of long actin filaments or bundles (S. Zhang et al. 2016; Suarez et al. 2015; Paul and Pollard 2008). One study shows that the polymerization rate increases ∼2–3 fold in vitro when the barbed end of actin filaments is associated with a processive formin, AtFH14 (S. Zhang et al. 2016). Alternatively, a previous in vitro study shows a non-processive formin – *Arabidopsis* FORMIN1 (AtFH1) – associates with the side of a pre-existing filament to nucleate new filaments (Michelot et al. 2006). Overall, experimental studies suggest that Arp2/3 complex and non-processive formins are responsible for the side branch nucleation of actin filaments, whereas processive formins bind to free barbed ends to enhance polymerization (Courtemanche, 2018).

Interplays between actin and actin-binding proteins govern the stochastic dynamics behavior of individual filaments in the cortical actin array of plant cells. However, the steady-state actin array morphology can be altered during pathogen invasion, triggering the first layer of innate immunity in plant cells called pattern-triggered immunity (PTI). Pattern recognition receptors detect both microbe-associated molecular patterns (MAMPs) and damage-associated molecular patterns, initiating mitogen-activated protein kinase and calcium-dependent protein kinase cascades (Macho & Zipfel, 2014; Ngou et al., 2021; Wang et al., 2022). During PTI, several cellular processes also occur, including fluctuations in cytosolic calcium concentration, reactive oxygen species production, accumulation of phospholipids, and cytoskeletal remodeling (B. Li et al. 2016; L. Cao et al. 2022; Torres, Dangl, and Jones 2002; Keinath et al. 2015; De Jong et al. 2004; Henty-Ridilla et al. 2013). Previous in vivo studies demonstrate that the dynamics of single actin filaments are altered during PTI, with MAMPs inducing an increase in actin filament density, length, and lifetime (Henty-Ridilla et al. 2013; Henty-Ridilla et al. 2014). Further studies reveal a significant reduction in severing and capping activity in epidermal plant cells during PTI, due to inhibition of ADF/cofilin and capping protein, respectively, contribute to the increased filament abundance (L. Cao et al., 2022; Henty-Ridilla et al., 2014; J. Li, Henty-Ridilla, et al., 2015; J. Li et al., 2017).

Our recent study investigated whether Arp2/3 complex and formins cooperate to generate and maintain the homeostatic actin array in plant epidermal cells and found that, while the individual inhibition of either Arp2/3 complex or formins reduced actin filament abundance, simultaneous inhibition of both nucleators surprisingly increased the frequency of the de novo nucleation of actin filaments, leading to an overall increase in actin filament abundance (Xu et al., 2024). Although the exact mechanism underlying the increased nucleation of actin filaments remains unclear, the availability of monomers likely plays a key role. We speculate that this may be due to the signal-mediated inhibition of a profilin isoform, thereby releasing monomeric actin for de novo nucleation (Xu et al., 2024).

Computational studies can provide valuable insights into understanding the dynamics and underlying mechanism of individual actin filament dynamics, especially when the coordinated action of a myriad of different actin-binding proteins modulates filament behaviors. Although several computational models have been developed for the actin cytoskeleton in mammalian cells, there have been only a few models specifically designed to simulate the cortical actin array in plant cells. A few existing models focus primarily on specific subprocesses within the plant actin cytoskeleton. For example, one model was developed for cytoplasmic streaming with consideration of actin bundles and myosin motors to describe how different patterns of streaming are obtained (Woodhouse & Goldstein, 2013). Recent studies employed coarse-grained simulations of actin filaments and cross-linkers to identify morphometric parameters that characterize the synthetic actin array, including density, orientation, ordering, and bundling (Akenuwa et al., 2024; Akenuwa & Abel, 2023). Although these approaches were able to reproduce the bundling dynamics of actin filaments, they did not consider individual actin filament dynamics in detail. In this study, we propose an agent-based model that aims to simulate the single filament dynamics in the homeostatic cortical actin array of plant epidermal cells. Our computational model was able to reproduce steady-state conditions in terms of the density, length, and population of actin filaments as well as the rates of severing, polymerization, and depolymerization.

## RESULTS

### Stochastic dynamics of actin filaments in plant epidermal cells

In epidermal cells from the dark grown hypocotyl, the cortical array comprises a sparsely populated, randomly distributed collection of actin filament bundles and single filaments (Smertenko et al., 2010; Staiger et al., 2009). These subpopulations are distinguishable based on fluorescence intensity, lifetime, and dynamic behavior (Staiger et al., 2009). New single filaments originate or are nucleated either de novo in the cytoplasm (Fig. 1A: Supplemental Movie 1) or as side branches on pre-existing filaments or bundles (Fig. 1B; Supplemental Movie 2). Approximately 8 nucleation events are observed in a 400-µm^2^ area during 100-s timelapse imaging, or 2×10^-4^ events/µm^2^/s (Xu et al. 2024; Table 1). In the homeostatic array, side-branched filament nucleation is 50% more frequent than de novo nucleation and occurs at the rate of 1.2×10^-4^ events/µm^2^/s, whereas de novo nucleation occurs at 0.8×10^-4^ events/µm^2^/s (Xu et al., 2024). Growth at the filament barbed end is rapid and occurs at rates between 1.25 µm/s and > 3 µm/s, with an overall average of 1.6–1.8 µm/s (Fig. 1C; Supplemental Movie 3; Table 1). By reducing the activity of Arp2/3 complex or formins genetically or with chemical inhibition coupled with quantitative analysis of the frequency distributions of different subpopulations of growing filaments, it is inferred that free barbed ends elongate at a rate of ∼1.25 µm/s, whereas processive formins accelerate growth to 2–2.25 µm/s (Xu et al., 2024). The pointed ends of filaments, if observed to shrink at all, depolymerize at a rate that is an order of magnitude slower or ∼0.2 µm/s (Fig. 1D; Supplemental Movie 4; Table 1). Thus, individual filaments do not display treadmilling behavior characterized by a balance between the growth rate at barbed ends and the shrinkage rate at pointed ends. Filaments grow to an average length of 10–17 µm before growth ceases, presumably through barbed-end capping (Fig. 1E; Supplemental Movie 5), followed by disassembly through prolific severing activity (Fig. 1F; Supplemental Movie 6). Individual filaments have an average lifetime of 15–20 s. Filament severing occurs at the rate of 0.008–0.014 events/µm/s (Table 1). This overall dynamic behavior of filament nucleation, rapid growth, and disassembly through severing and slow depolymerization is referred to as stochastic dynamics, and produces a dynamic steady state that is distinct from treadmilling (J. Li, Blanchoin, et al., 2015; Staiger et al., 2009).

**Figure 1.**
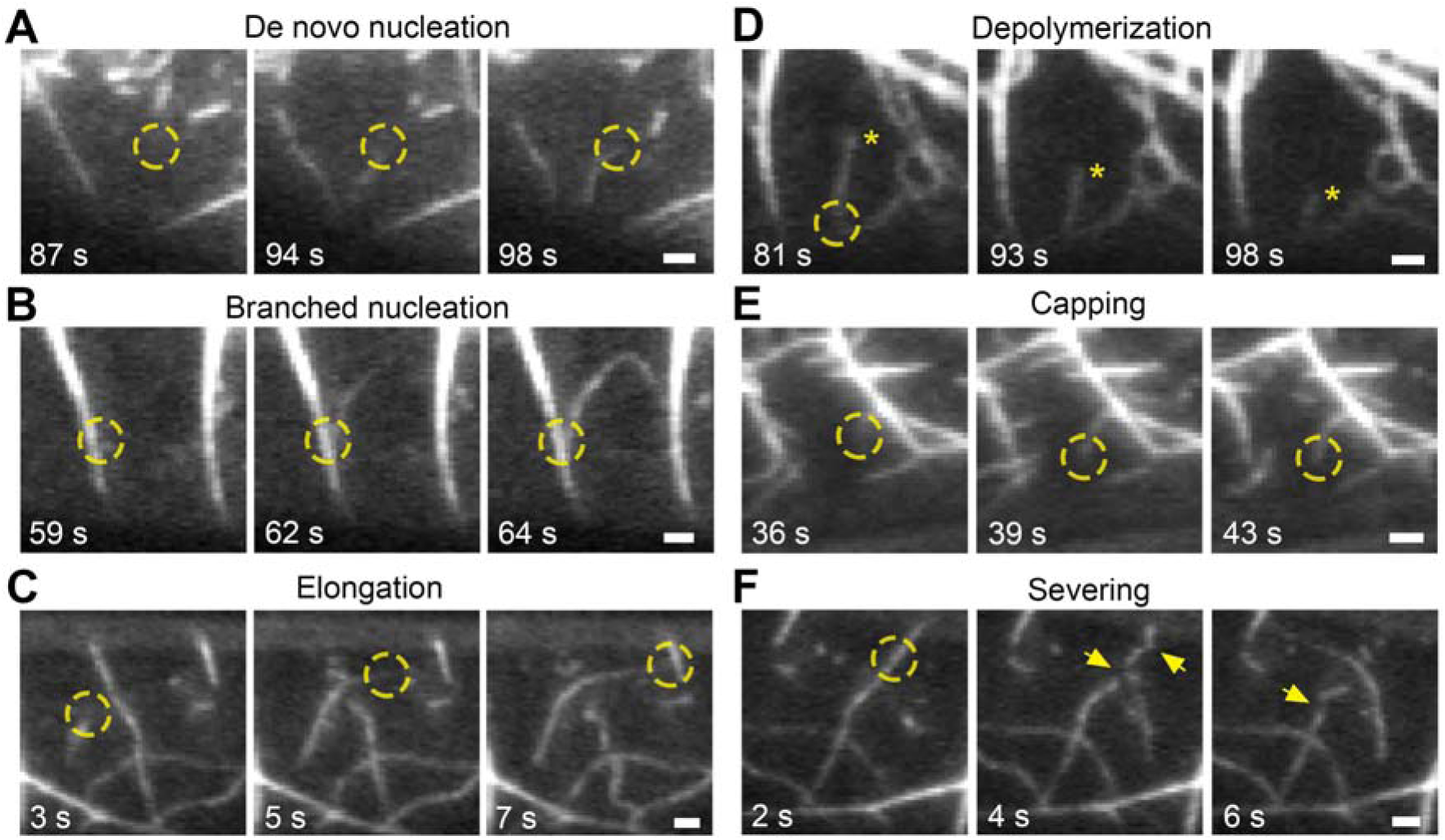
Single actin filament dynamics of the cortical array in Arabidopsis hypocotyl epidermal cells are revealed by live-cell imaging. Representative time series of cortical actin filament dynamics collected with variable-angle epifluorescence microscopy of dark-grown hypocotyl epidermal cells expressing the GFP-fABD2 reporter. (A, B) New filaments initiated either de novo in the cytoplasm (A) or from the side of an existing filament or bundle (B). (C) An example of actin filament elongation/polymerization occurring at a rate of ∼2 µm/s. (D) Depolymerization of a filament from the pointed end (yellow asterisk) at a slow rate, ∼0.2 µm/s. (E) Filament growth from a barbed end ceased, suggesting a capping event. (F) A filament is disassembled by multiple severing events (yellow arrows). Yellow circles indicate the sites of nucleation, elongation, capping, and severing of actin filaments. Scale bars indicate 2 µm.

**Table 1.** Quantities measured from simulations, compared with experimentally determined values. Experimental values were obtained from (L. Cao et al., 2016; Chaudhry et al., 2007; Henty-Ridilla, Li, et al., 2013; Staiger et al., 2009; Xu et al., 2024; S. Zhang et al., 2016). The nucleation frequency was recalculated as events/µm^2^/s, using data from (Xu et al., 2024).

| Stochastic dynamic parameters | Experiment | Model |
| --- | --- | --- |
| Polymerization rate [$\mu\text{m}/\text{s}$] | 1.6–1.8 | $2.2 \pm 0.4$ |
| Depolymerization rate [$\mu\text{m}/\text{s}$] | 0.17–0.20 | $0.2 \pm 0.01$ |
| Enhancement of polymerization by formin | 2–3 | 3 |
| Filament length [$\mu\text{m}$] | 10–17 | $9.9 \pm 0.85$ |
| Severing frequency [events/$\mu\text{m}/\text{s}$] | 0.008–0.014 | $0.01 \pm 0.002$ |
| Nucleation frequency [events/$\mu\text{m}^2/\text{s}$] | $2 \times 10^{-4}$ | $1.20 \times 10^{-4}$ |
| Fraction of profilin-bound actin monomers | 0.99 | 0.99 |
| Fraction of profilin-free actin monomers | 0.01 | 0.01 |

### An agent-based model reproduces the stochastic filament dynamics

We developed an agent-based computational model that simulates the stochastic dynamic behavior of individual actin filaments, incorporating key molecular interactions observed in vivo to achieve a dynamic steady state. In our model, actin filaments are simplified into serially connected cylindrical segments with polarity defined by barbed and pointed ends (Fig. 2A). Because differences between branches formed by Arp2/3 complex and those formed by non-processive formin remain unclear in terms of association and dissociation rates and geometrical configurations, we include only the Arp2/3 complex in our model. The Arp2/3 complex is simplified into two cylindrical segments connected at its center point. The positions of all the segments are updated via the Langevin equation based on Brownian dynamics.

**Figure 2.**
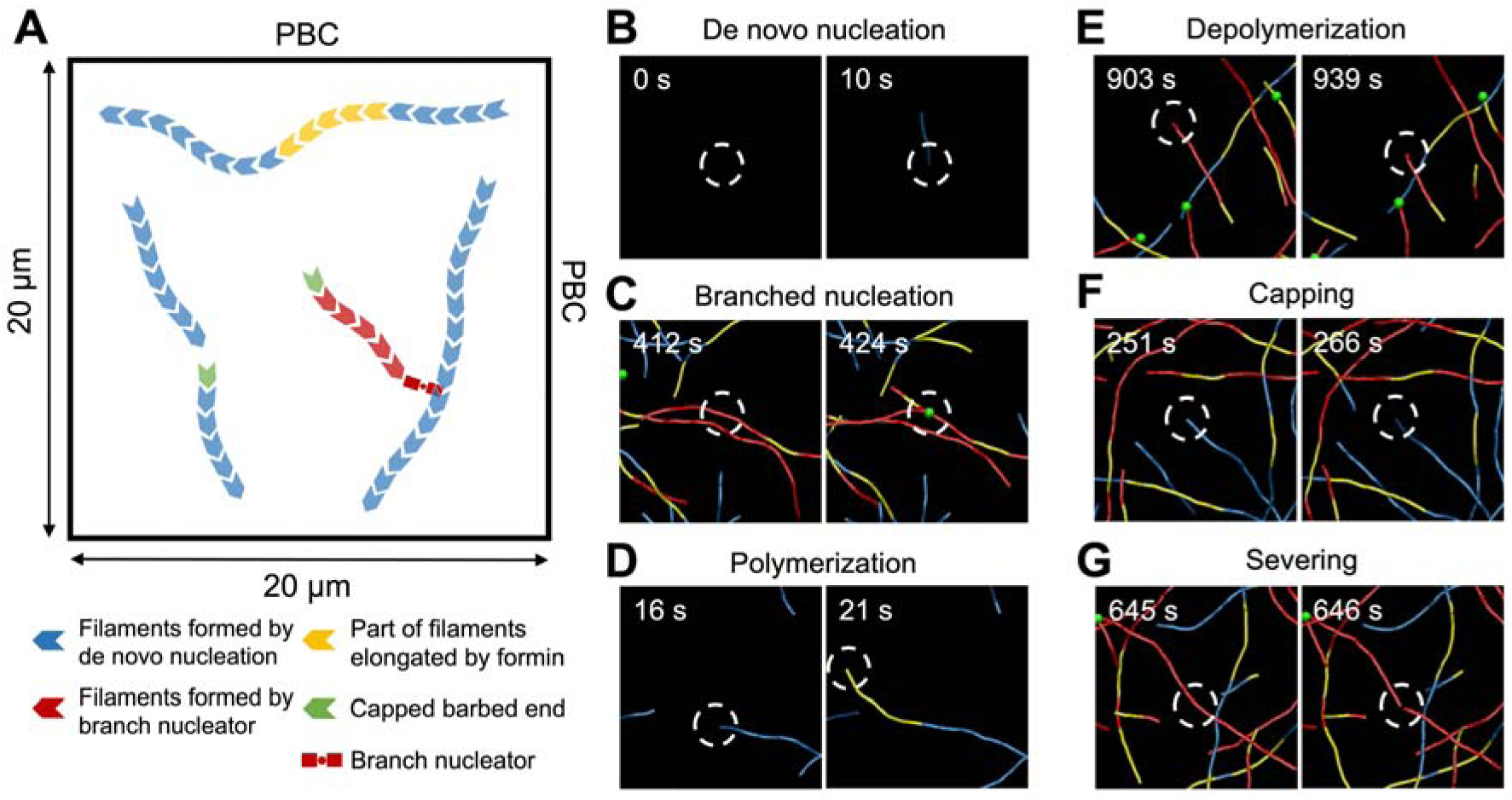
Agent-based computational model of the homeostatic cortical actin array in plant epidermal cells. (A) Model schematic depicting actin filament dynamics. Filaments originate via de novo nucleation (blue) or branched nucleation (red). Part of these filaments elongated by 3-fold faster formin-mediated polymerization is marked in yellow. Capped barbed ends (green) are unable to polymerize further. Actin segments in filaments transition from the ATP state to the ADP state and are susceptible to severing and depolymerization. Simulations were performed in a 20×20×1 µm domain with periodic boundary conditions (PBC) applied along the x- and y-axes. (B–G) Snapshots showing key dynamic behaviors occurring in the model. (B) De novo nucleation: actin filaments emerge spontaneously. (C) Branched nucleation: the binding of a branched nucleator to an existing filament results in a branched nucleation event. (D) Polymerization: the barbed end of a filament elongates slowly via spontaneous polymerization or faster with formin-mediated polymerization (yellow). (E) Depolymerization: the pointed end of a filament in the ADP state shrinks over time. (F) Capping: free barbed ends transition to the capped state, preventing further elongation. (G) Severing: part of the actin filaments in the ADP state undergo severing, promoting filament disassembly. White circles indicate the sites of nucleation, polymerization, depolymerization, capping, and severing of actin filaments.

To mimic the situation in vivo, actin filaments undergo de novo nucleation, branched nucleation, polymerization, depolymerization, capping, and severing (Fig. 2A). The de novo nucleation occurs using only profilin-free actin monomers (Fig. 2B). The Arp2/3 complex binds to the side of existing filaments to nucleate a new filament (Fig. 2C). The barbed end of actin filaments can transition between a free state to a formin-bound state, which exhibits 3-fold faster polymerization using only profilin-bound actin monomers (Fig. 2D). The pointed end of actin filaments in the ADP state is prone to depolymerization (Fig. 2E). The barbed end can also transition between the free state and a capped state. The barbed end in the capped state is not allowed to elongate via polymerization (Fig. 2F). Uncapping of the barbed end can also occur, but we set its rate extremely low relative to the capping rate to reflect in vivo conditions where most of the capped filaments eventually disassemble (J. Li et al., 2012; Staiger et al., 2009). Based on a prior experimental study (McCullough et al., 2011), we assume that filament severing occurs in an angle-dependent manner (Fig. 2G). As before, the severing rate is determined by two parameters: zero-angle severing rate constant (*k*_0,sev_) and angle sensitivity (λ_sev_) (Eq. 9). A balance between actin assembly and disassembly allows the actin array to reach a dynamic steady state.

Using the model, we explored a wide parametric space to identify conditions that reproduce the steady state of the cortical actin array in plant epidermal cells (Table 2). We categorized actin filaments in the actin array into three classes. Under the reference condition (Table 2), the relative fraction of the three classes showed a steady state after ∼500 s (Fig. 3A; Supplemental Movie 7): branched filaments (i.e., filaments formed by Arp2/3 complex) elongated by slow polymerization (∼50%), de novo filaments (i.e., filaments formed by the de novo nucleation) elongated by slow polymerization (∼20%), and filaments elongated by formin-enhanced polymerization (∼30%) (Fig. 3B). Accordingly, at steady state, our model produced a considerably higher number of branched filaments than de novo filaments within a 400-µm^2^ area (Fig. 3C). The nucleation frequency, calculated as the total number of de novo and branched nucleation events divided by the domain size (i.e., size of xy plane) and total simulation time, was 1.2×10C^4^ events/µm^2^/s. Branched nucleation occurred twice as frequently as de novo nucleation, with a rate of 0.8×10C^4^ events/µm^2^/s, whereas de novo nucleation occurred at a rate of 0.4×10C^4^ events/µm^2^/s. Average filament length was 9.3 ± 1.1 µm for de novo filaments and 9.1 ± 0.4 µm for branched filaments (Fig. 3D). The average polymerization rate was measured to be 2.2 ± 0.4 µm/s, whereas the depolymerization rate was 0.2 ± 0.01 µm/s (Fig. 3E). The severing rate was measured to be 0.009 ± 0.002 events/µm/s (Fig. 3F), and the number of capped barbed ends was 42.1 ± 6.3 (Fig. 3G). Note that we measured these parameters in units previously used for biological experiments (J. Li & Staiger, 2018; Xu et al., 2024), to allow direct comparison between simulations and experiments (Table 1). Values from 1000 s onward were used for computation to focus on the steady-state results. We found that the simulation values were typically within 2-fold or less of the empirically determined values (Table 1). In the following simulations, we varied each of the key parameters to understand how the actin array is regulated.

**Figure 3.**
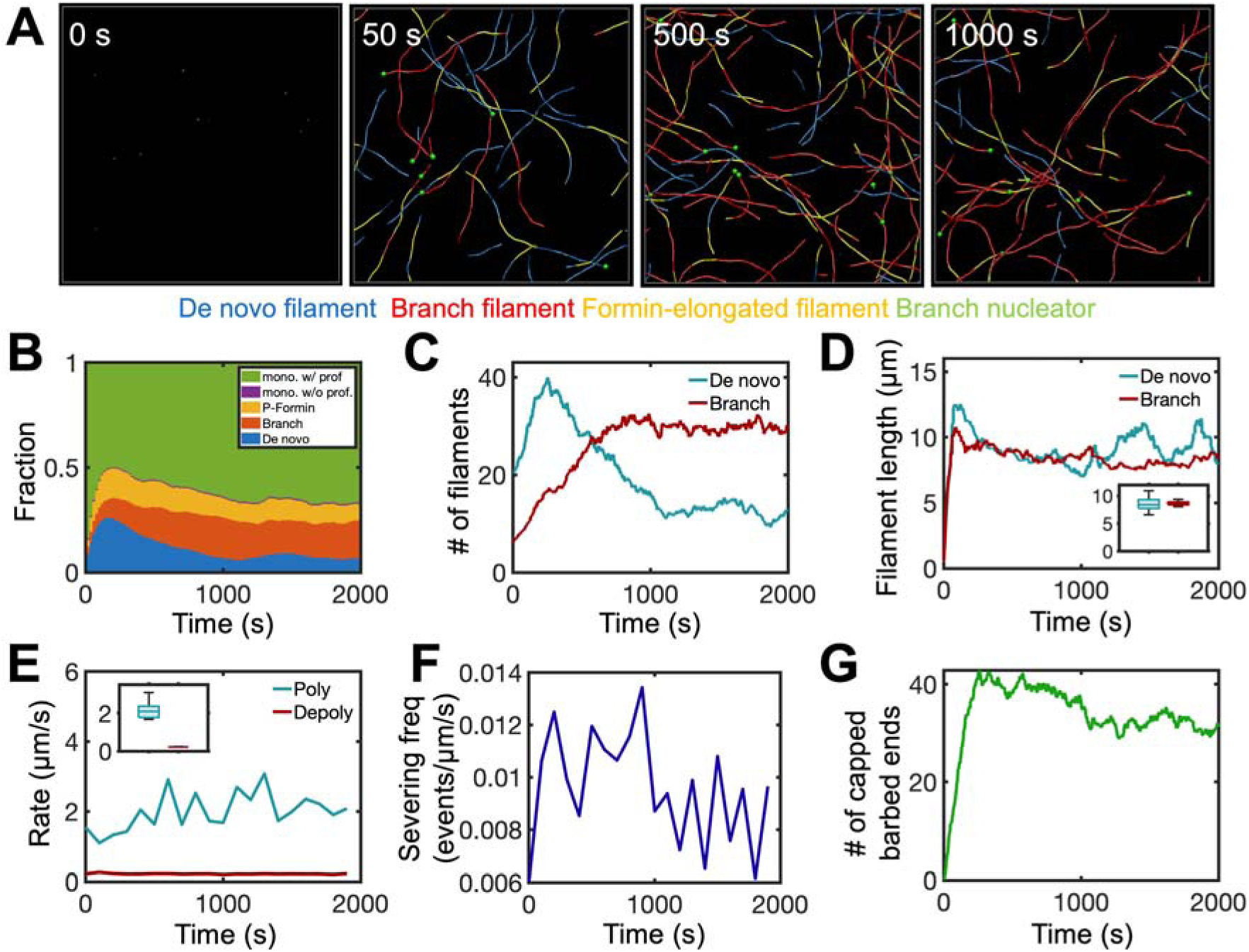
Steady-state dynamics of the cortical actin array in plant epidermal cells under the reference condition. (A) Snapshots of an actin network at 0, 50, 500, and 1000 s under the reference condition, showing the spatiotemporal organization of filaments formed via de novo nucleation (blue) and branched filaments (red) formed via branch nucleators (green). Part of these filaments elongated faster by formin and are represented by yellow. (B) The fraction of actin in 5 different states over time: profilin-bound monomer (green), profilin-free monomer (purple), formin-elongated filaments (yellow), branched filaments elongated slowly (red), and filaments formed by de novo nucleation and slow polymerization (blue). (C) Time course of the number of de novo (blue) and branched (red) filaments. (D) Time course of the average length for de novo (blue) and branched (red) filaments. Inset: average filament length of de novo versus branched filaments. (E) Polymerization (blue) and depolymerization (red) rates measured throughout the simulation. Inset: average rates for polymerization versus depolymerization. (F) Time course of severing frequency. (G) Time course of the number of capped barbed ends. In the insets (D, E), boxes show interquartile range with a horizontal line indicating the median. Whiskers extend from the box to the minimum and maximum data values. Data were averaged from five independent simulations, and values from 1000 s onward were used in the box plots to focus on the steady-state results.

**Table 2.**
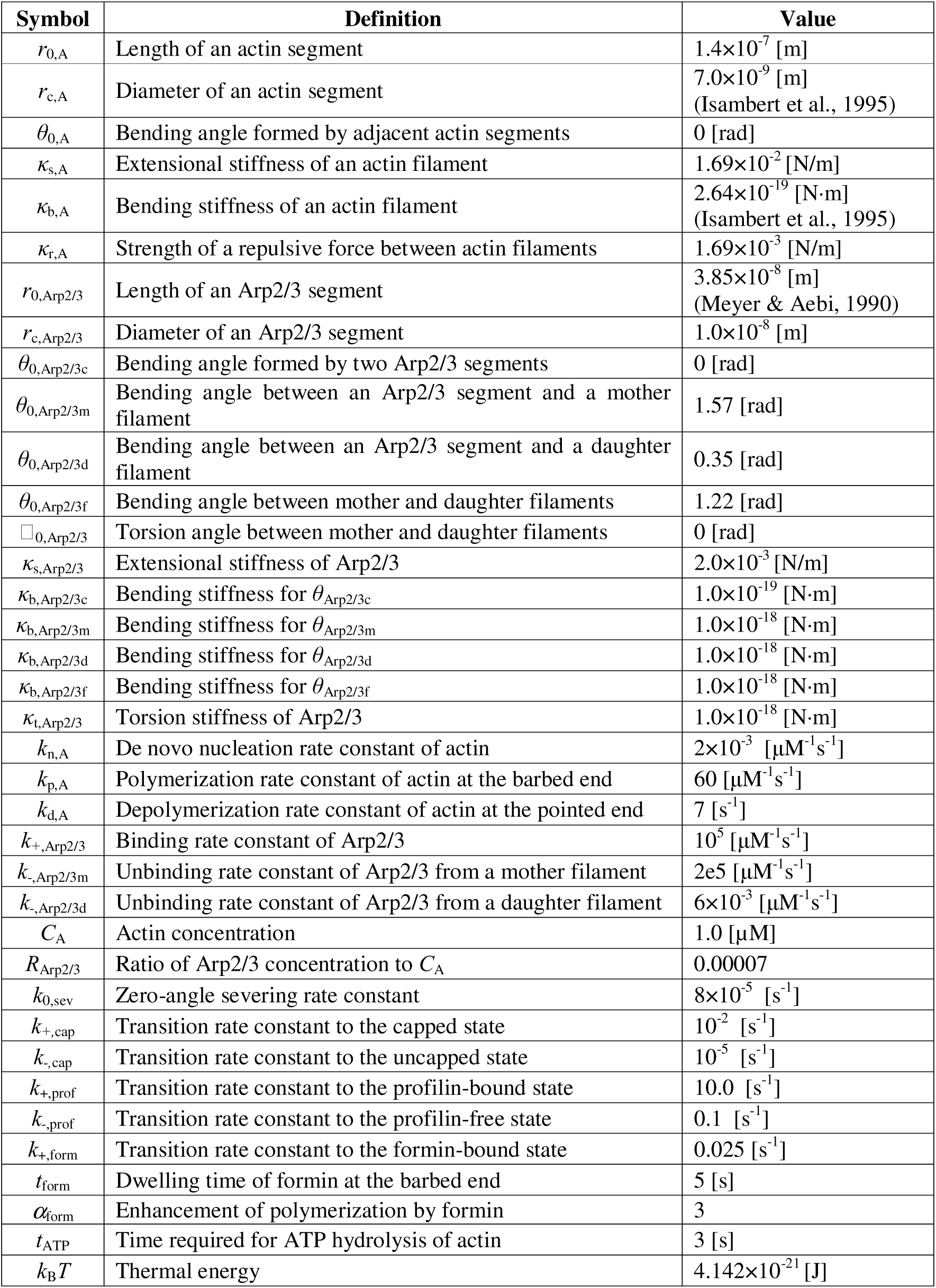
List of parameters used in the plant cytoskeleton model. Each parameter was varied over a wide range to choose the reference condition, with some values derived from previous studies.

### Inhibition of severing and capping disrupts the steady state of the actin array

Given that the average polymerization rate at the barbed end is ∼10-fold higher than the depolymerization rate at the pointed end in hypocotyl epidermal cells (Staiger et al., 2009), the enhancement of filament turnover through severing and capping activities is crucial for maintaining the dynamic steady state (Henty et al., 2011; J. Li et al., 2012). We investigated how severing and capping maintain the steady state by enhancing or inhibiting these activities. First, we probed the effects of varying severing rate. As expected, increasing the severing rate led to a greater number of shorter filaments overall and more frequent severing events (Figs. 4A, B). The numbers of both de novo and branched filaments increased with the higher severing rate (Figs. 4C, D), whereas the length of both types of filaments decreased (Figs. 4E, F). Severing results in enhanced filament disassembly by creating more pointed ends for depolymerization and capping newly formed barbed ends rather than merely dividing each filament into two. It was indeed observed that the fraction of filaments in the actin pool decreased with an increasing severing rate (Fig. 4G). Due to the elevated fraction of monomers at high severing activity, de novo nucleation frequency also increased—rising progressively from 2.0×10C^5^ to 4.2×10C^5^ events/µm^2^/s as *k*_0,sev_ increased from 0 to 24×10C^5^ s^-1^. These results are consistent with the changes in the number and turnover of actin filaments resulting from genetic mutation of a single ADF/cofilin isoform (Henty et al., 2011; Henty-Ridilla et al., 2014)

**Figure 4.**
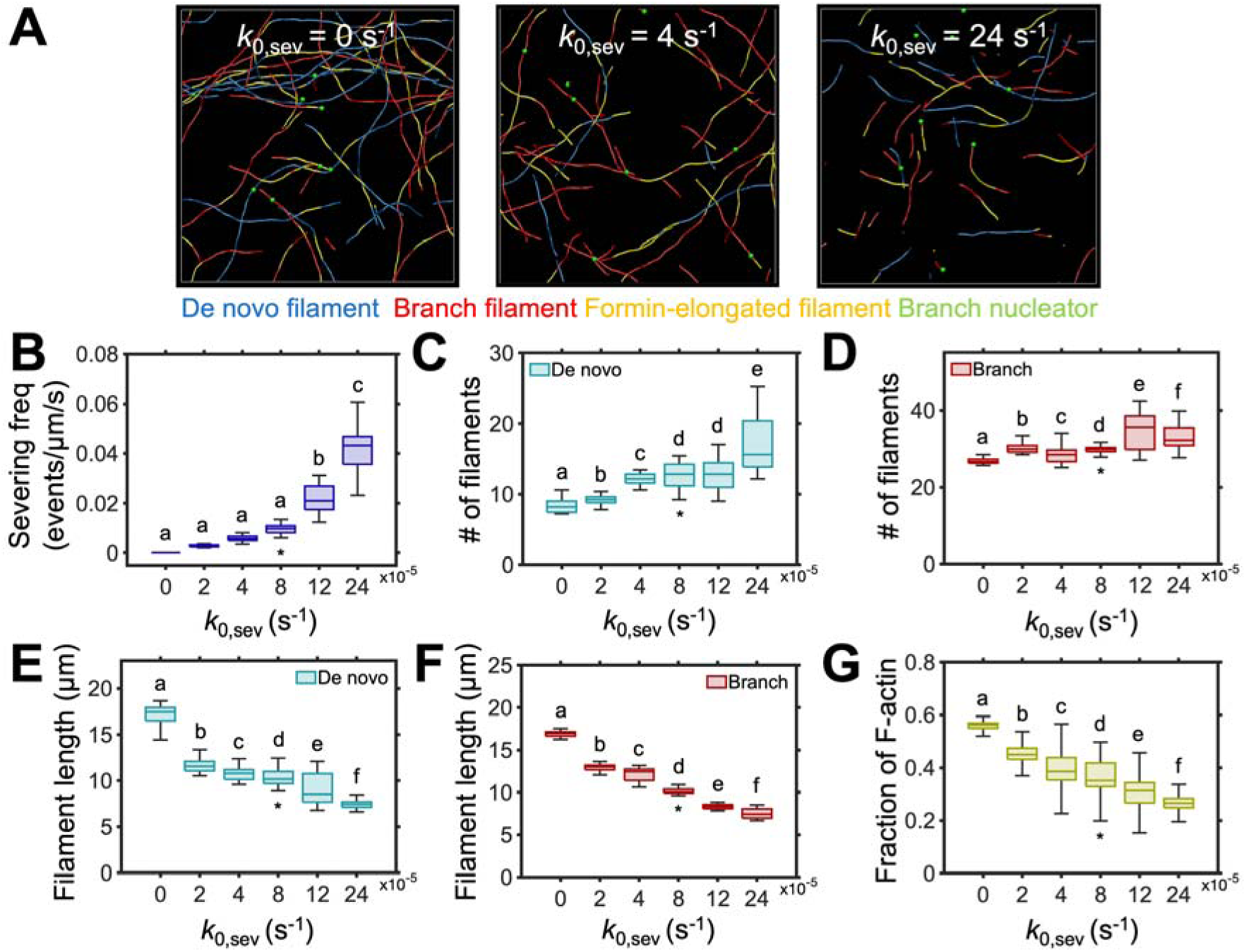
Severing increases actin filament abundance and reduces filament length. (A) Snapshots of actin networks taken at ∼1000 s generated with a different zero-angle severing rate constant (*k*_0,sev_): 0 s^-1^ (no severing), 4 s^-1^ (moderate severing), and 24 s^-1^ (frequent severing). (B) The frequency of severing events increases with higher *k*_0,sev_. (C, D) The number of de novo and branched filaments with different *k*_0,sev_, showing that severing increases the total number of filaments. (E, F) The average length of de novo and branched filaments depends on *k*_0,sev_. Filaments become shorter with more frequent severing. (G) The fraction of actin in the filamentous state as a function of *k*_0,sev_. A decrease in this fraction suggests that severing enhances filament disassembly, leading to an increased concentration of the actin monomer pool. In panels (B–G), boxes show interquartile range with a horizontal line indicating the median. Whiskers extend from the box to the minimum and maximum data values. Data were averaged from five independent simulations per condition, and values from 1000 s onward were used in the box plots to focus on the steady-state results. Letters indicate groups that are significantly different based on one-way ANOVA with Tukey’s post-hoc test (p < 0.05). Asterisks (*) indicate the reference condition.

Next, we varied the capping rate to assess its contribution to the abundance and turnover of actin filaments. As the capping rate increased, a larger fraction of the barbed ends became capped, resulting in a much sparser actin array (Figs. 5A, B). Consequently, the numbers of both de novo and branched filaments decreased (Figs. 5C, D). Excessive capping severely restricted actin assembly, leading to shorter filaments (Figs. 5E, F). Similar to the case with the excessive severing rate, the high capping rate resulted in a decrease in the fraction of filaments in the actin pool due to excessive F-actin disassembly (Fig. 5G). The elevated fraction of monomers at high capping activity led to an increase in de novo nucleation frequency—rising progressively from 0.4×10C^5^ to 6.6×10C^5^ events/µm^2^/s as *k*_+,cap_ increased from 0 to 0.04 s^-1^. These results are consistent with changes in the number and turnover of actin filaments resulting from genetically-induced variation in the heterodimeric capping protein levels (Li et al. 2012; 2014; 2015).

**Figure 5.**
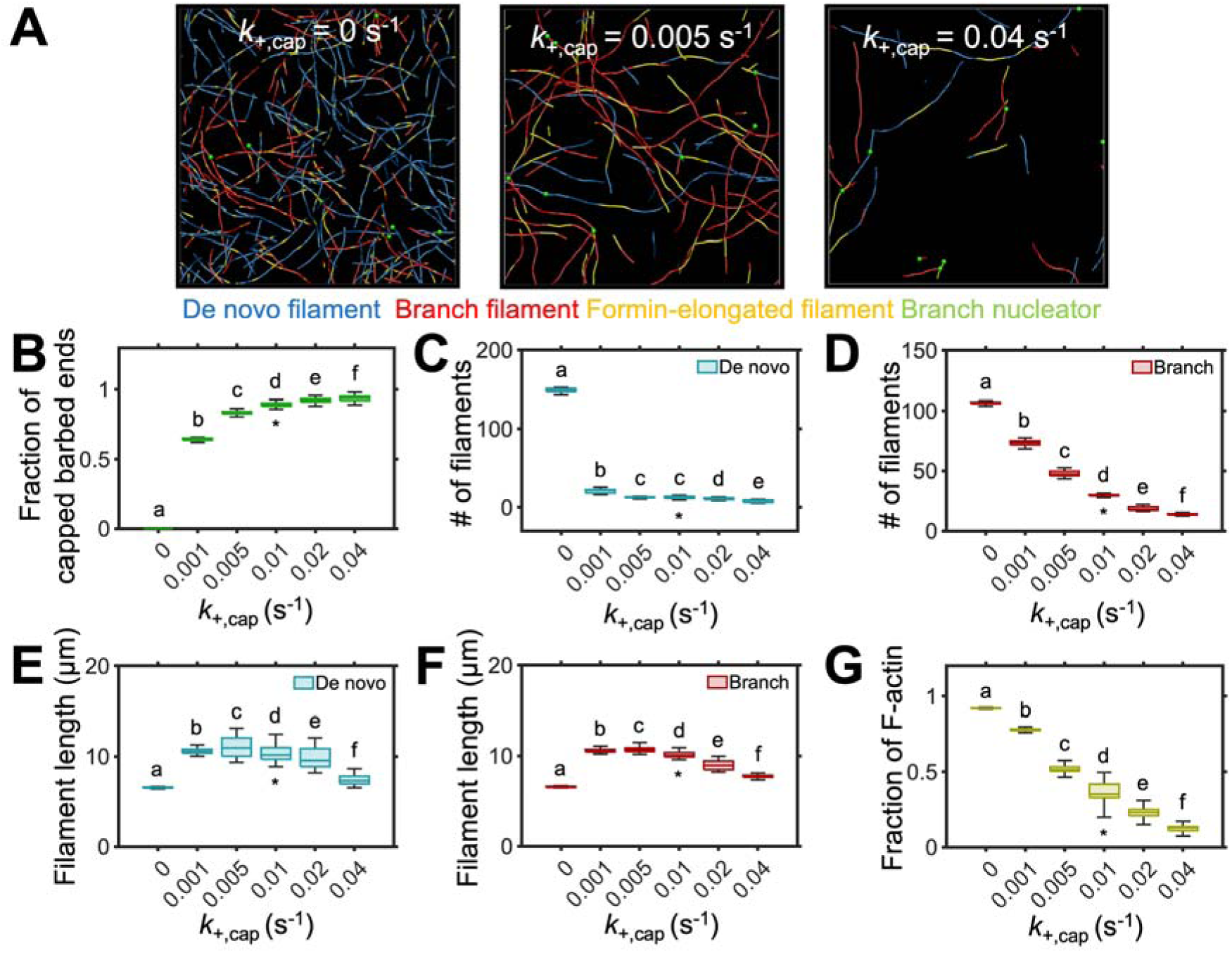
Capping reduces the abundance and length of actin filaments. (A) Snapshots of actin networks taken at ∼1000 s generated with a varying transition rate from the free state to the capped state (*k*_+,cap_): 0 s^-1^ (no capping), 0.005 s^-1^ (moderate capping), and 0.04 s^-1^ (frequent capping). The capped barbed ends cannot elongate further. (B) The fraction of capped barbed ends increases with higher *k*_+,cap_. (C, D) The number of de novo and branched filaments with different *k*_+,cap_, showing a substantial decrease in total filament abundance with the increased capping rate. (E, F) The average length of de novo and branched filaments was reduced as capping activity was enhanced. Without capping, a large number of short filaments were present, reducing the average filament length. (G) The average fraction of actin in the filamentous state with different *k*_+,cap_. The decline in F-actin fraction indicates that capping promotes filament disassembly, leading to a larger actin monomer pool. In panels (B–G), boxes show interquartile range with a horizontal line indicating the median. Whiskers extend from the box to the minimum and maximum data values. Data were averaged from five independent simulations per condition, and values from 1000 s onward were used in the box plots to focus on the steady-state results. Letters indicate groups that are significantly different based on one-way ANOVA with Tukey’s post-hoc test (p < 0.05). Asterisks (*) indicate the reference condition.

During the plant defense response, MAMP treatments elicit a rapid increase in filament abundance, presumably through the inhibition of the severing and capping of actin filaments (Henty-Ridilla et al., 2014; J. Li, Henty-Ridilla, et al., 2015). When we inhibited both capping and severing in the agent-based model, the fraction of actin in the filamentous form and the density of actin filaments increased markedly compared to the reference case (Figs. 6A, B). Without the two disassembly-promoting mechanisms, the number of filaments continued to increase (Fig. 6C), as the polymerization rate at the barbed end was always faster than the depolymerization rate at the pointed end. As a result, filament length also increased compared to the reference case (Fig. 6D). Notably, with the actin concentration of 1 µM, the actin pool was depleted rapidly (Fig. 6B), which became a limiting factor for further actin assembly (Fig. 6E). Under this condition, the de novo nucleation frequency decreased to 0.4×10C^5^ events/µm²/s, representing a 10-fold reduction compared to the reference case. Considering that the concentration of free actin monomers is much higher in plant cells (Chaudhry et al. 2007), both the density and length of filaments observed in this simulation are likely to be the underestimation of experimental measurements. These results demonstrate that severing and capping are essential for proper disassembly and turnover of F-actin, and they are crucial for maintaining actin filament homeostasis. The results also reaffirm that ADF4 and CP are key targets for inhibition by defense response signaling that lead to actin remodeling and increased filament abundance (Henty-Ridilla et al., 2014; J. Li, Henty-Ridilla, et al., 2015).

**Figure 6.**
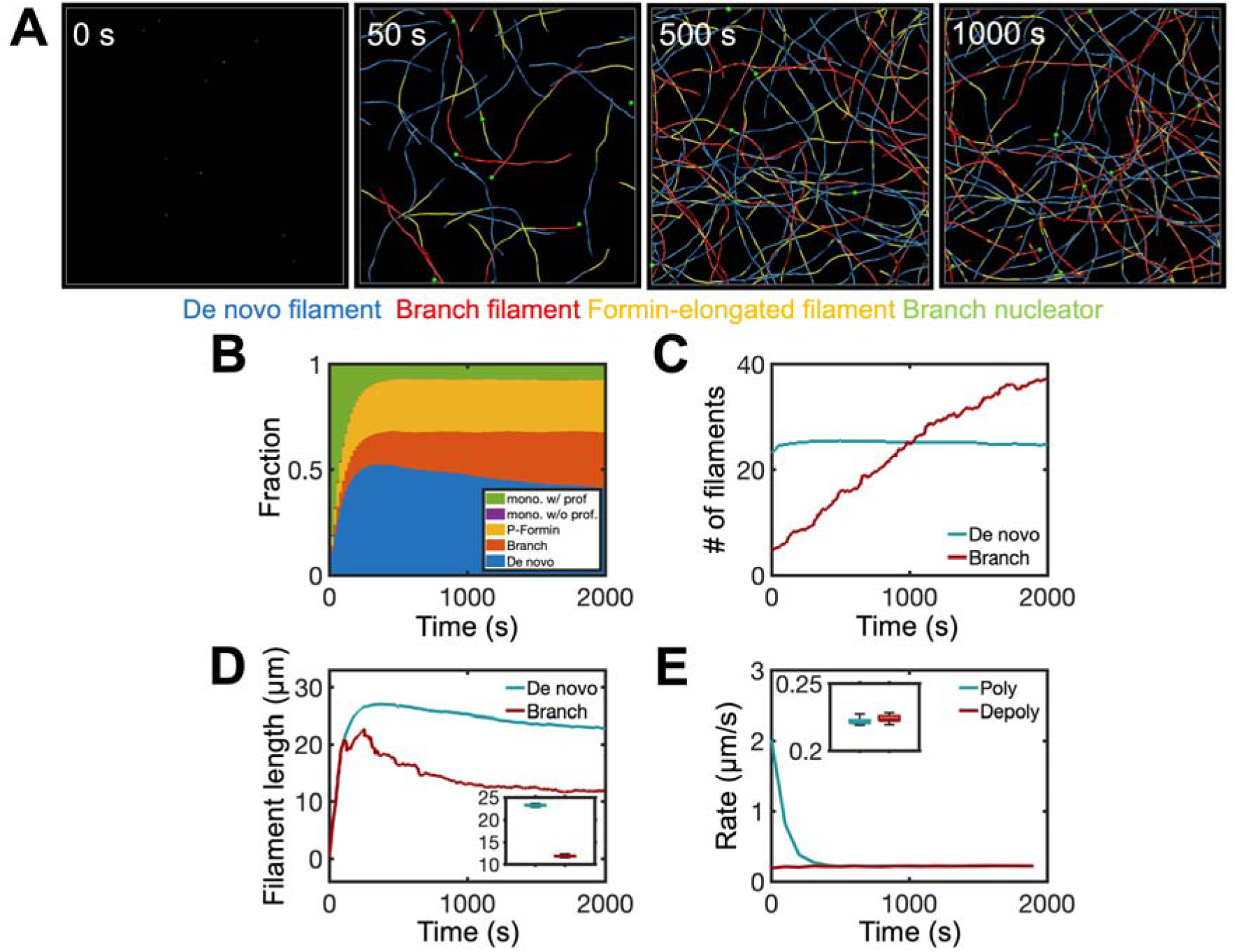
Simultaneous inhibition of severing and capping increases actin filament abundance and length. (A) Snapshots of an actin network taken at 0, 50, 500, and 1000 s with both severing and capping inhibited. This simulation recapitulated the state of the actin array in plant epidermal cells observed during pattern-triggered immunity. (B) The fraction of actin in different states over time, showing a substantial increase in F-actin density due to the inhibition of severing and capping. (C) Time evolution of the number of de novo (blue) and branched (red) filaments, indicating the continuous accumulation of branched filaments. (D) Time evolution of the average length of de novo (blue) and branched (red) filaments. Inset: filament length distribution. (E) Polymerization (blue) and depolymerization (red) rates measured over 2000 s. Inset: rate distribution. The depletion of the actin pool seen in (A) became a limiting factor for filament polymerization. After monomers were depleted, the polymerization rate was almost identical to the depolymerization rate because polymerization can take place only after monomers are available from depolymerization. In the insets (D, E), boxes show interquartile range with a horizontal line indicating the median. Whiskers extend from the box to the minimum and maximum data values. Data were averaged from five independent simulations, and values from 1000 s onward were used in the box plots to focus on the steady-state results.

### Branched-filament nucleation and formin-enhanced polymerization are crucial for achieving the steady state of the actin array

Branched-filament nucleation plays a crucial role in the formation of actin arrays by generating new filaments along existing filaments, which compensates for slow de novo nucleation (Xu et al., 2024). When we decreased Arp2/3 density (*R*_Arp2/3_), the fraction of branched filaments in the array noticeably decreased (Figs. 7A, B). While the number of de novo filaments modestly increased, the number of branched filaments significantly decreased at lower Arp2/3 densities (Figs. 7C, D). However, the average length of both filament types remained largely unchanged compared to the reference case (Figs. 7E, F). The overall fraction of actin in the filamentous state decreased with reduced branched nucleation resulting from lower Arp2/3 density (Fig. 7G). As *R*_Arp2/3_ decreased from 9×10C^5^ to 0, the increased monomer pool led to an increase in de novo nucleation frequency from 3.2×10C^5^ to 5.0×10C^5^ events/µm^2^/s.

**Figure 7.**
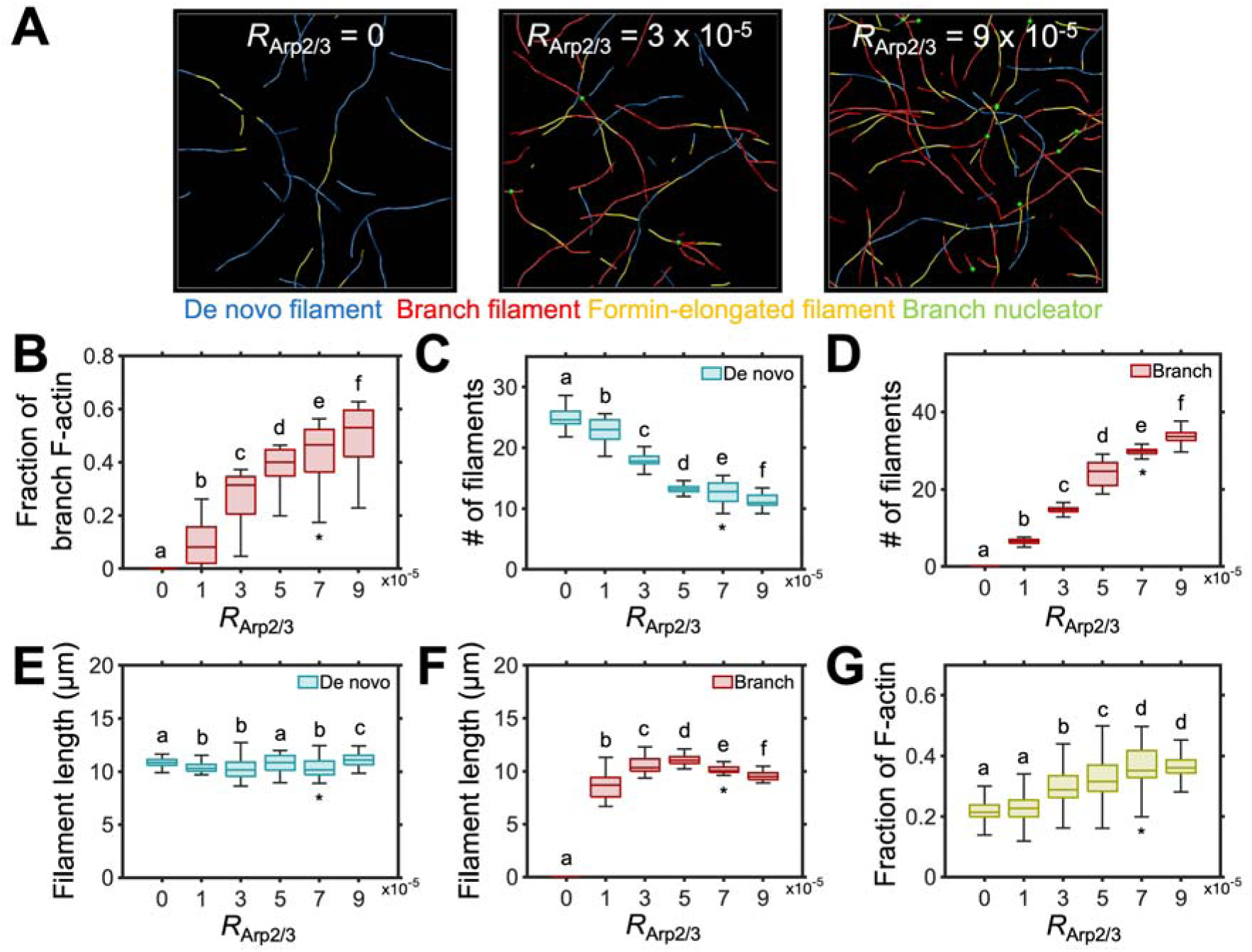
Adequate Arp2/3 complex density is essential for establishing the steady-state actin network architecture. (A) Snapshots of actin networks taken at ∼1000 s with varying Arp2/3 densities (*R*_Arp2/3_): 0, 3×10LJ^5^, and 9×10LJ^5^. (B) The fraction of branched filaments decreases with lower *R*_Arp2/3_, indicating reduced nucleation activity. (C, D) The number of de novo and branched filaments as a function of *R*_Arp2/3_, showing a substantial decrease in the number of branched filaments with lower *R*_Arp2/3_, whereas the number of de novo filaments increased slightly. (E, F) The average length of de novo and branched filaments. (G) The fraction of actin in the filamentous state decreases with lower Arp2/3 density, reflecting reduced branch nucleation. In panels (B–G), boxes show interquartile range with a horizontal line indicating the median. Whiskers extend from the box to the minimum and maximum data values. Data were averaged from five independent simulations per condition, and values from 1000 s onward were used in the box plots to focus on the steady-state results. Letters indicate groups that are significantly different based on one-way ANOVA with Tukey’s post-hoc test (p < 0.05). Asterisks (*) indicate the reference condition.

We also probed the effect of processive formin by varying the transition rate from the free state to the formin-bound state (*k*_+,form_). Similar to the effect observed with reducing Arp2/3 density, a lower transition rate led to a decreased fraction of actin filaments elongated by formin (Fig. 8A). The polymerization rate of actin filaments also decreased with lower transition rates, reflecting the loss of an enhanced elongation rate associated with processive formin-associated barbed ends (Fig. 8B). The number and length of both de novo and branched filaments showed a modest decrease or remained relatively stable with lower transition rates, which could be due to the reduction of filament lifetime from slower elongation rate (Figs. 8C–F). Consequently, the fraction of actin in filamentous state showed a modest decrease with lower transition rates (Fig. 8G). As *k*_+,form_ decreased from 0.105 to 0 s^-1^, the increased monomer pool was accompanied by an increase in de novo nucleation frequency from 2.1×10C^5^ to 4.4×10C^5^ events/µm^2^/s.

**Figure 8.**
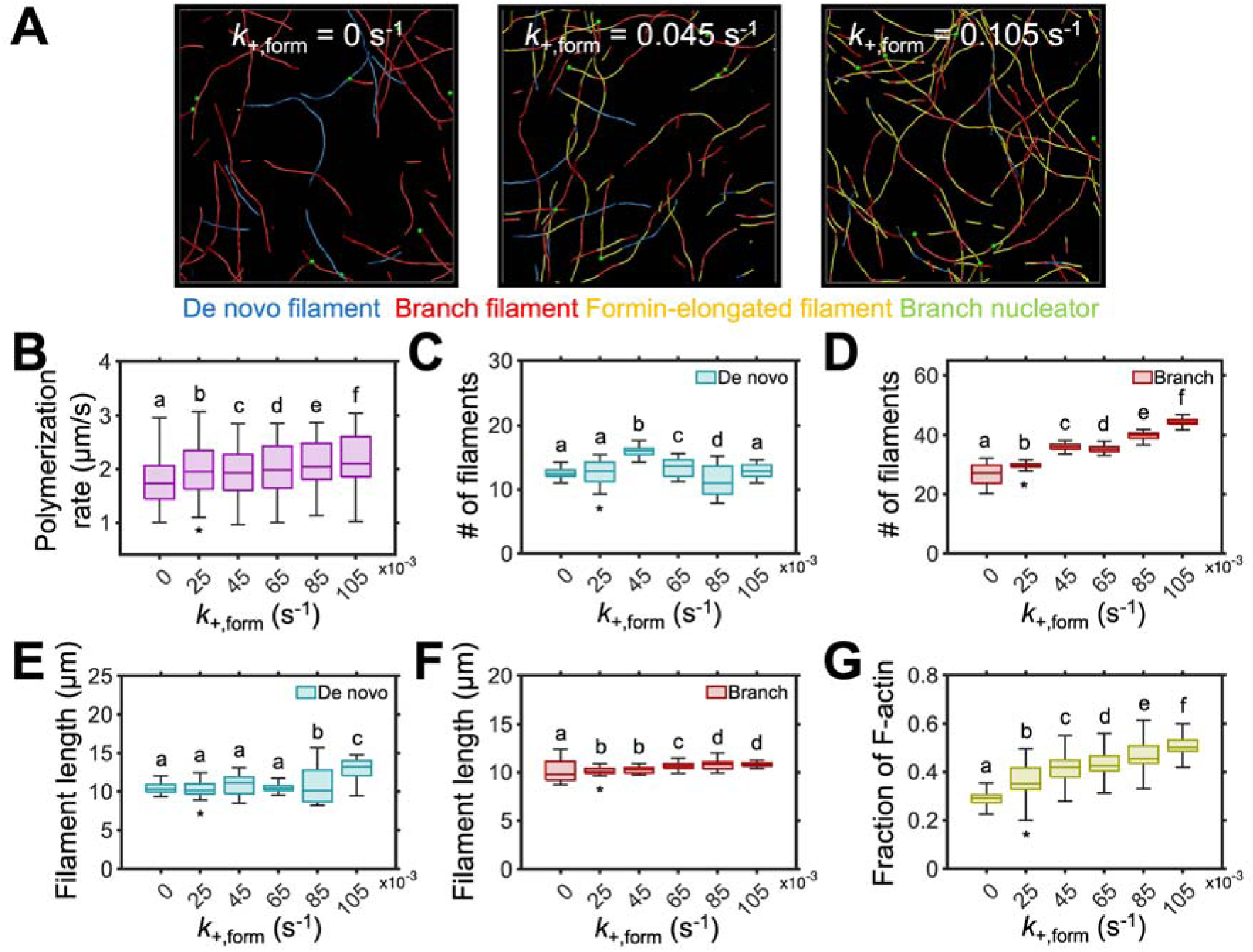
Enhanced processive formin activity increases the actin polymerization rate. (A) Snapshots of actin networks taken at ∼1000 s with a varying formin transition rate (*k*_+,form_): 0 s^-1^, 0.045 s^-1^, and 0.105 s^-1^. *k*_D,formin_ represents a transition rate from the free state to the formin-bound state at the barbed end, which results in a 3-fold increase in the polymerization rate. (B) Actin polymerization rates increase with elevated formin activity, highlighting the role of processive elongation in actin array dynamics. (C, D) The number of de novo and branched filaments with different *k*_+,form_, showing minimal changes in filament population. (E, F) The average length of de novo and branched filaments, which initially increased with *k*_+,form_, but plateaued at higher values as indicated by overlapping statistical groups. (G) The average fraction of actin in the filamentous state increases with higher *k*_+,form_. In panels (B–G), boxes show interquartile range with a horizontal line indicating the median. Whiskers extend from the box to the minimum and maximum data values. Data were averaged from five independent simulations per condition, and values from 1000 s onward were used in the box plots to focus on the steady-state results. Letters indicate groups that are significantly different based on one-way ANOVA with Tukey’s post-hoc test (p < 0.05). Asterisks (*) indicate the reference condition.

Based on our recent finding that simultaneous genetic and/or chemical inhibition of Arp2/3 complex and formins increases actin filament abundance through a 2.5-fold enhanced de novo nucleation frequency (Xu et al., 2024), we next probed the effect of inhibiting both the branched-filament nucleator and processive formin activity. We found that the overall actin array morphology was disrupted, with a decrease in F-actin abundance as de novo nucleation became the sole contributor to the actin array formation (Figs. 9A, B). Under this condition, the de novo nucleation frequency increased to 0.53×10C^4^ events/µm²/s, which is a ∼33% increase compared to the reference case. The number of de novo filaments increased by 44.9% relative to the reference case, whereas their length showed minimal change (Figs. 9C, D). Additionally, the rates of polymerization, depolymerization, and severing frequency did not change noticeably compared to the reference case (Figs. 9E–F). Following the decrease in F-actin density, the number of capped barbed ends also declined when both Arp2/3 and formins were inhibited (Fig. 9G). Collectively, these results demonstrate that the Arp2/3 complex and formins act cooperatively to contribute to the dynamic steady state of the cortical actin array and are necessary for achieving the filament density observed in vivo.

**Figure 9.**
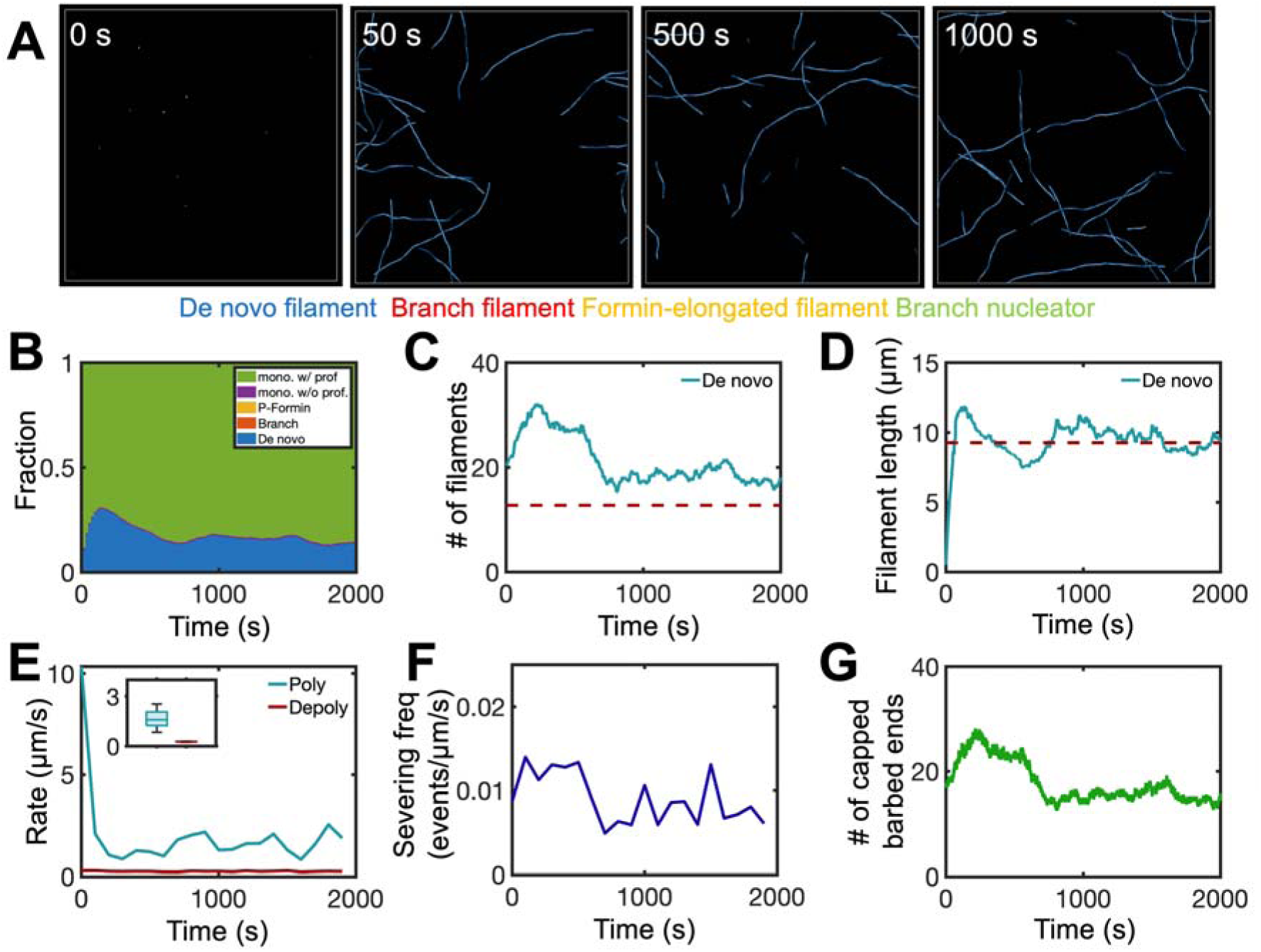
Simultaneous inhibition of Arp2/3 complex and processive formin activity leads to a reduction in overall filament density. (A) Snapshots of an actin network taken at 0, 50, 500, and 1000 s with both Arp2/3 complex and formin inhibited. (B) The fraction of actin in different states over time, showing a marked decrease in filament density due to the absence of branched nucleation and formin-mediated assembly. (C, D) Time course of the number and average length of de novo filaments. Red dashed line indicates the steady-state count and length of de novo filaments from the reference case. (E) Polymerization (blue) and depolymerization (red) rates over 2000 s. Inset: rate distribution. (F) Severing frequency remains unchanged relative to the reference case. (G) The number of capped barbed ends declines as the total filament count decreases due to the absence of Arp2/3 and formin. In the inset (E), boxes show interquartile range with a horizontal line indicating the median. Whiskers extend from the box to the minimum and maximum data values. Data were averaged from five independent simulations, and values from 1000 s onward were used in the box plots to focus on the steady-state results.

### Regulation of profilin is likely responsible for the enhanced de novo nucleation when both Arp2/3 complex and formins are inhibited

Our model produced a drastically different result compared to experimental studies, which showed a marked increase in de novo filament nucleation when Arp2/3 complexes and formins were simultaneously inhibited in plant epidermal cells (Xu et al., 2024). This is likely due to an increase in free monomer concentration, perhaps resulting from signal-mediated down regulation of profilin activity generating more actin monomers for spontaneous nucleation. We tested whether increasing the transition rate for actin monomers from a profilin-bound state to a profilin-free state (*k*_-,prof_) could reproduce the experimental findings. Similar to the conditions in Fig. 9, both Arp2/3 complex and formin activities were set to zero in this case (Figs. 10A). Increasing the transition rate reduced the fraction of the profilin-bound monomers in the actin pool and conversely increased the availability of profilin-free monomers for de novo nucleation (Figs. 10B, C). As *k*_-,prof_ increased from 0.1 to 10 s^-1^, the de novo nucleation frequency rose correspondingly from 4.8×10C^5^ to 5.7×10C^3^ events/µm²/s. As the number of de novo filaments increased substantially (Fig. 10C), the free actin monomer pool was depleted and the fraction of actin in the filamentous state increased (Figs. 10E). As described earlier, this depletion of the free actin monomer pool became a limiting factor, restricting the filament length (Fig. 10D).

**Figure 10.**
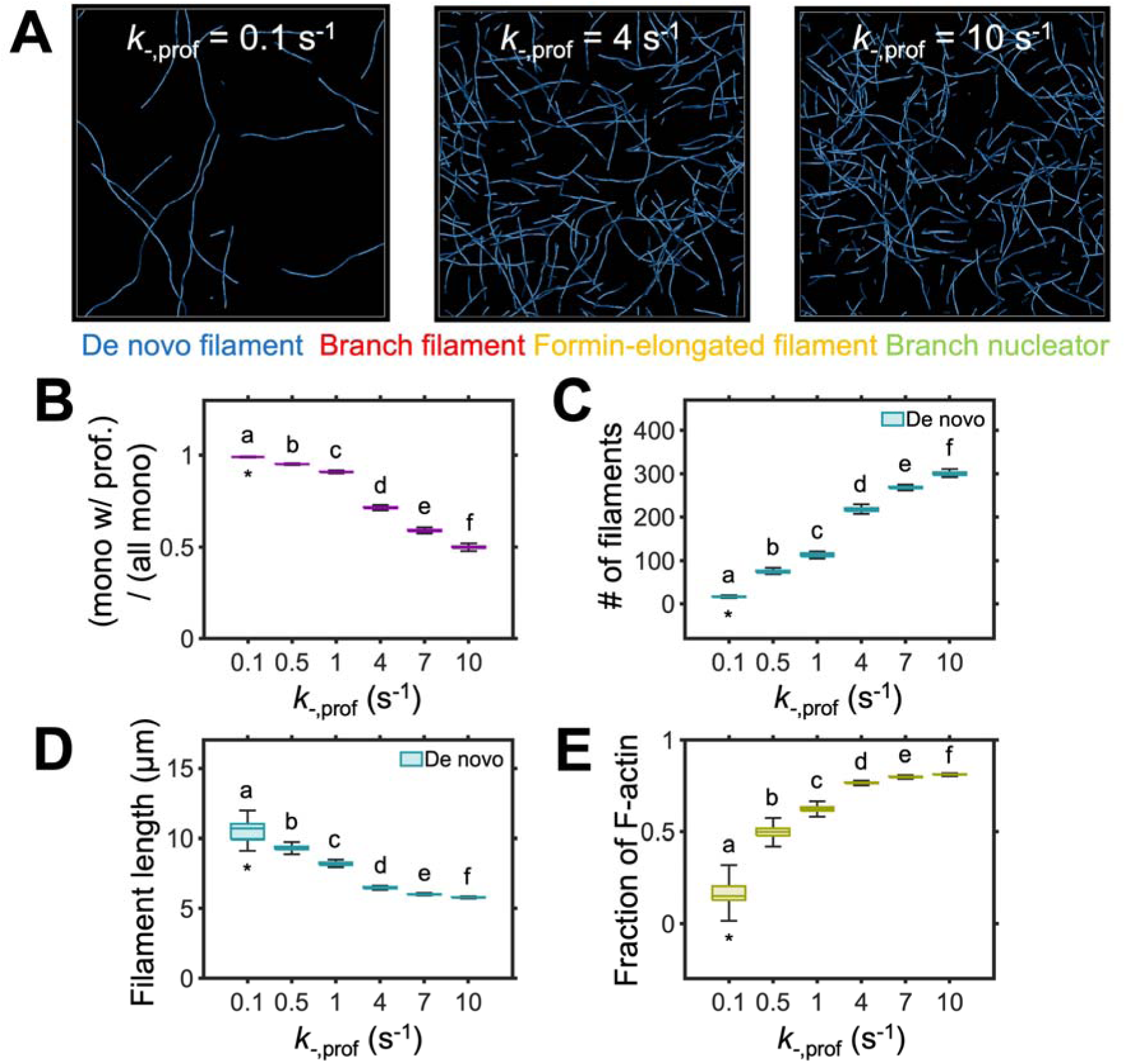
Suppression of profilin activity and the inhibition of both Arp2/3 complex and formin enhance de novo nucleation, increasing overall filament density. (A) Snapshots of actin networks taken at ∼1000 s with a varying profilin transition rate (*k*_-,prof_): 0.1 s^-1^, 4 s^-1^, and 10 s^-1^. *k*_-,prof_ represents a transition rate from the profilin-bound state to free state of actin monomers. (B) The fraction of monomers with profilin in the actin pool with different *k*_-,prof_. (C, D) The number and average length of de novo filaments depending on *k*_-,prof_, showing an increase in abundance and a decrease in length. (E) The fraction of actin in the filamentous state. Lower *k*_-,prof_ decreases the concentration of profilin-free monomers, which suppresses de novo nucleation and leads to a significant reduction in filament density. In panels (B–E), boxes show interquartile range with a horizontal line indicating the median. Whiskers extend from the box to the minimum and maximum data values. Data were averaged from five independent simulations per condition, and values from 1000 s onward were used in the box plots to focus on the steady-state results. Letters indicate groups that are significantly different based on one-way ANOVA with Tukey’s post-hoc test (p < 0.05). Asterisks (*) indicate the reference *k*_-,prof_ value.

We repeated these simulations in the presence of both Arp2/3 and formin activities (Fig. 11). The observation was overall similar, but the number and length of branched filaments decreased with a higher transition rate as de novo nucleation dominated the array. As *k*_-,prof_ increased from 0.1 to 10 s^-1^, the de novo nucleation frequency rose correspondingly from 0.4×10C^4^ (reference case) to 4.2×10C^3^ events/µm²/s. These results highlight profilin as a key regulator of de novo nucleation, with the availability of profilin-free actin monomers playing a crucial role in filament assembly. Our findings help reconcile discrepancies between simulations and experimental observations when the activity of both nucleator classes are suppressed and predict the importance of profilin regulation in actin array organization.

**Figure 11.**
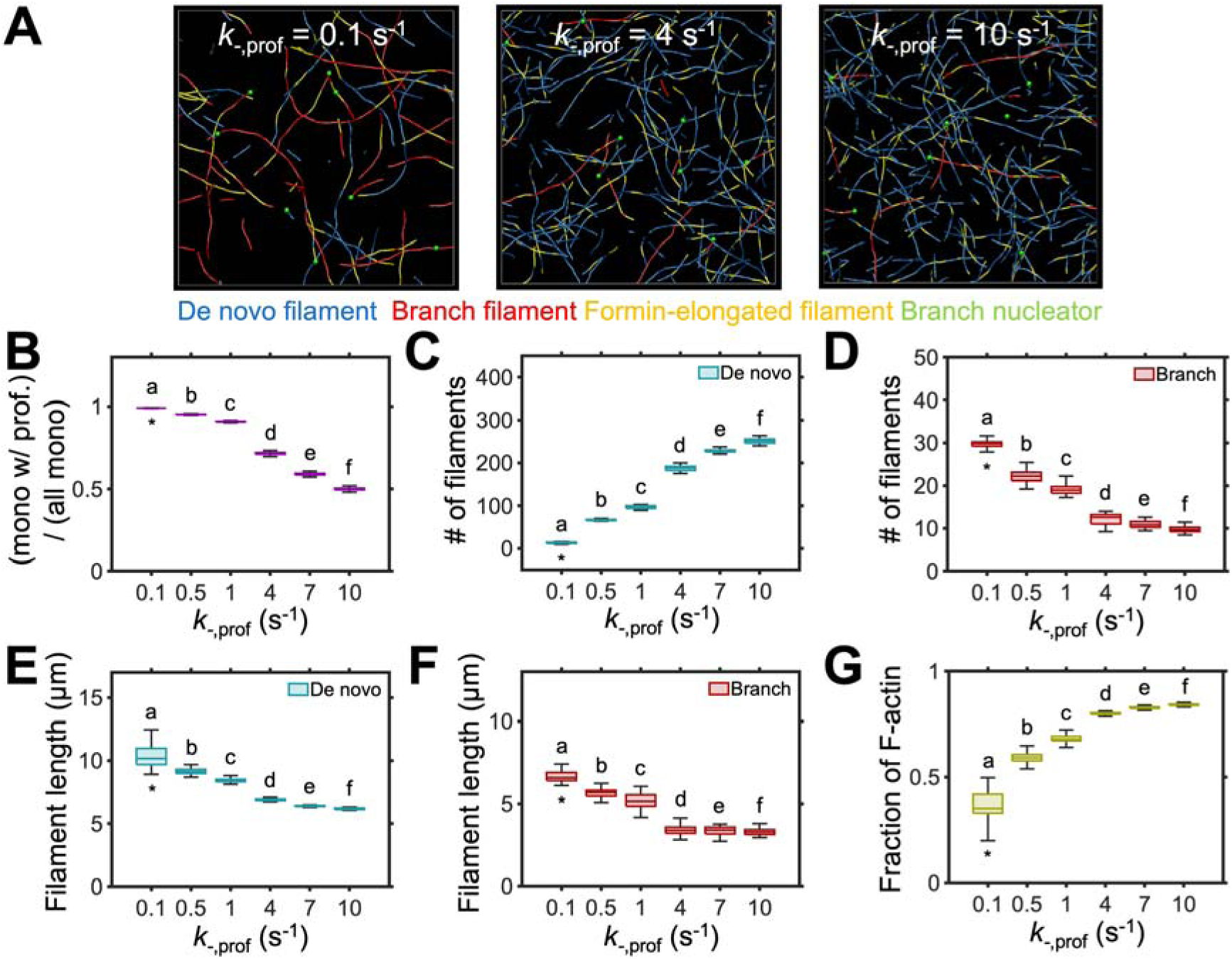
Suppression of profilin activity promotes de novo filament formation, allowing it to dominate in the actin array architecture. (A) Snapshots of representative actin networks at ∼1000 s generated with a different profilin transition rate (*k*_-,prof_): 0.1 s^-1^, 4 s^-1^, and 10 s^-1^. (B) The fraction of monomers bound to profilin in the actin pool decreases with higher *k*_-,prof_. (C, D) The number of de novo and branched filaments with different *k*_-,prof_, showing a substantial increase in the number of de novo filaments and a modest decrease in the number of branched filaments as *k*_-,prof_ increases. (E, F) The average length of de novo and branched filaments. (G) The fraction of actin in the filamentous state. With lower *k*_-,prof_, de novo nucleation was suppressed due to an increased proportion of profilin-bound actin monomers, which increases the availability of actin monomer pool for filament elongation. In panels (B–G), boxes show interquartile range with a horizontal line indicating the median. Whiskers extend from the box to the minimum and maximum data values. Data were averaged from five independent simulations per condition, and values from 1000 s onward were used in the box plots to focus on the steady-state results. Letters indicate groups that are significantly different based on one-way ANOVA with Tukey’s post-hoc test (p < 0.05). Asterisks (*) indicate the reference condition.

## DISCUSSION

In this study, we developed an agent-based model to simulate single actin filament dynamics in the homeostatic cortical actin array of plant epidermal cells. In Arabidopsis epidermal cells, the cortical array comprises a sparse network of intermingled actin filaments and bundles, exhibiting a dynamic steady-state behavior characterized by de novo nucleation, branched nucleation, rapid polymerization, slow depolymerization, capping, and prolific severing activities (Fig. 1).

Using our model, we simulated the effects of various molecular factors to capture the stochastic dynamics of individual filaments observed in vivo (Fig. 2). We successfully reproduced key features of the homeostatic cortical actin array, including the abundance and length of actin filaments and the rate or frequency of severing, polymerization, and depolymerization (Fig. 3). Through a comprehensive parametric study, we highlighted the importance of severing and capping activities in maintaining actin turnover and disassembly and were able to recapitulate findings from genetic loss- and/or gain-of-function studies with CP and/or ADF mutants. Consistent with experimental studies, excessive severing resulted in a larger number of short filaments (Fig. 4), whereas the inhibition of severing led to fewer, longer filaments (Henty et al., 2011; Henty-Ridilla et al., 2014; Zheng et al., 2013). Conversely, excessive capping restricted filament elongation and reduced both the density and length of filaments (Fig. 5), whereas inhibiting capping led to a substantial increase in filament density (J. Li et al., 2017; J. Li, Henty-Ridilla, et al., 2015).

We then simulated a state representative of plant epidermal cells during PTI, where both severing and capping activities are suppressed by signal-mediated cytoplasmic fluxes in second messengers (Henty-Ridilla et al., 2014; J. Li, Henty-Ridilla, et al., 2015; Wang et al., 2022). Consistent with experimental observations, inhibiting both capping and severing noticeably increased filament abundance and length compared to the reference case, which rapidly depleted the actin monomer pool (Fig. 6). These results emphasize the critical role of severing and capping in filament turnover in the homeostatic cortical array, which is particularly essential given the rapid elongation rate of actin filaments in epidermal cells.

In our model, actin filaments can be generated from de novo nucleation, branched nucleation, or processive formins associated with pre-existing barbed ends. Supporting experimental findings (Cifrová et al., 2020; Xu et al., 2024), we found that decreasing the density of the branched nucleator resulted in the inhibition of branched filaments in the array morphology (Fig. 7). By contrast, an increase in the binding rate of processive formins led to a larger fraction of filaments elongated from pre-existing ends (Fig. 8), which induced a higher polymerization rate (S. Zhang et al. 2016). Despite these changes, the overall filament length did not vary noticeably, perhaps due to the prolific severing and capping activities. These findings demonstrate that both Arp2/3 complex and formins are essential for the formation of actin arrays in the homeostatic plant cortex.

Previous studies in animal cells have shown that inhibiting both the Arp2/3 complex and formins causes a marked decrease in the dynamics and turnover of the cortical actin array (Fritzsche et al., 2016). By contrast, our recent study on *Arabidopsis* epidermal cells showed an increase in filament abundance following genetic or chemical inhibition of both Arp2/3 and formins (Xu et al., 2024). Furthermore, we found the enhancement of de novo nucleation frequency as a sole contributor to this increased filament density. However, our modeling revealed that inhibiting both the Arp2/3 complex and formin resulted in a decrease in the total number of filaments, whereas the abundance and length of filaments remained unchanged (Fig. 9). One potential mechanism for enhanced de novo nucleation is through an increase in the pool of free monomers; the likeliest mechanism would be through down-regulation of profilin activity. We tested this with our model by decreasing the transition rate of actin monomers from the profilin-free state to the profilin-bound state to increase the fraction of profilin-free monomers available for de novo nucleation (Fig. 10). With triple inhibition of Arp2/3, formins, and profilin, we observed a significant increase in filament number, as well as a 100-fold increase in de novo nucleation, consistent with our recent findings (Xu et al., 2024). These results suggest that variations in the profilin-actin interaction can enhance the availability of free monomers, potentially boosting de novo nucleation and filament abundance in the plant cortex. This adaptive mechanism may occur in the plant epidermal cells when other forms of nucleation are impaired, but the exact signaling pathway behind this compensation remains unknown.

Although our model was able to recapitulate the roles of the key parameters measured from experimental studies, the model still has limitations. As described earlier, we used a relatively low actin concentration of 1 µM for the sake of computational efficiency. This resulted in a rapid depletion of the free actin monomer pool, which became a limiting factor for elongation of existing filaments. In *Arabidopsis* epidermal cells, total actin concentration likely ranges from 40–50 µM, the ratio of G- to F-actin exceeds 10:1, and there is a three-fold excess of profilin to total actin (Blanchoin et al. 2010; Chaudhry et al. 2007; Staiger et al., 2009; Xu et al., 2024). Additionally, this study aims to simulate single filament dynamics, and therefore we did not probe the effects of actin crosslinking proteins (ACPs) or myosin XI on the homeostatic actin array morphology. ACPs—such as villins, LIM proteins, and fimbrins—connect individual actin filaments to form actin bundles via a “catch and zipper” mechanism (Thomas, 2012). The quantification of bundle fluorescence intensity indicates that these ACPs form actin bundles comprised of 13 ± 5 filaments (Smertenko et al., 2010). These actin bundles play an important role in cytoplasmic streaming, where myosin XI walks along actin bundles for intracellular transport of various cargos (Tominaga et al., 2013). Studies have shown that loss of myosin XI function results in reduced filament severing and increased bundling, leading to decreased actin network dynamics (Cai et al., 2014). Additionally, myosin XI is responsible for the generation of forces that induce filament buckling and contribute to actin array remodeling (Cai et al., 2014). The roles of the Arp2/3 complex and formins in bundle formation and organization, the significance of severing and capping in bundle disassembly, and the impact of myosin XI on steady-state actin array dynamics will be explored in future studies.

Our model offers predictive insight into how network remodeling may occur in vivo. One of the key predictions is that de novo nucleation frequency increases in response to elevated severing or capping activity due to an expanded pool of free monomers (Figs. 4, 5). Conversely, dual inhibition of severing and capping leads to filament accumulation and monomer depletion and should lead to a reduced de novo nucleation frequency (Fig. 6). Notably, the model suggests that de novo nucleation does not rise following simultaneous inhibition of Arp2/3 and formin activities, unless profilin activity is also reduced— highlighting a critical interplay between filament disassembly and monomer availability (Figs. 9–10). This framework predicts that profilin could potentially act as a key regulator of compensatory nucleation, and suggests that experimental quantification of nucleation rates under altered severing, capping, and profilin conditions would directly test these mechanisms to further validate the model.

## CONCLUSIONS

In conclusion, our agent-based model successfully reproduced single filament dynamics in the homeostatic cortical actin array of plant epidermal cells. Our results indicated that inhibiting severing and capping significantly stagnates F-actin disassembly, which results in an elevated fraction of F-actin and limits the free actin monomer pool for new nucleation events. We further demonstrated that the simultaneous inhibition of Arp2/3 and formins does not increase de novo nucleation frequency, as observed in vivo. However, when profilin binding to actin monomers was additionally inhibited, de novo nucleation frequency was significantly increased, which sheds light on our prior experimental findings that the frequency of de novo nucleation increased when other forms of nucleation were inhibited. In the near future, we will incorporate ACPs and myosin XI to investigate their effects on cortical actin array organization and dynamics.

## MATERIALS AND METHODS

### Model Overview

#### Brownian dynamics via the Langevin equation

In our agent-based model, cytoskeletal elements—actin filament and Arp2/3—are simplified into serially connected cylindrical segments. The number of segments in each actin filament varies depending on filament length. Arp2/3 comprises two cylindrical segments connected at its center point.

The velocities of the endpoints of all segments are updated every time step by the Langevin equation with inertia neglected:

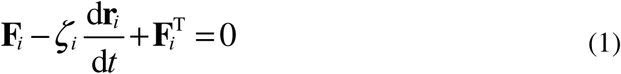

where **F***_i_*, ζ*_i_*, and **r***_i_* represent the deterministic force, drag coefficient, and position of the *i*th endpoint, respectively, and *t* is time. ζ*_i_* is calculated via an approximated form for a cylindrical object (Clift et al., 1978):

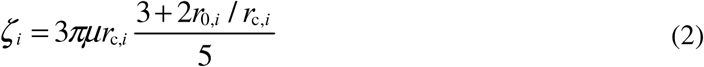

where *µ* is the viscosity of the surrounding medium, and *r*_0,*i*_ and *r*_c,*i*_ are the length and diameter of segments, respectively. **F***_i_*^T^ is a stochastic force satisfying the fluctuation-dissipation theorem (Underhill & Doyle, 2004):

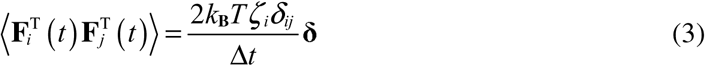

where δ is a second-order tensor, δ*_ij_* is the Kronecker delta, *k*_B_*T* is thermal energy, and Δ*t* = 1.15×10^-5^ s is time step. The positions of the cylindrical segments are updated every time step, using the velocities calculated via Eq. 1 (d**r***_i_*/d*t*) and the Euler integration scheme:

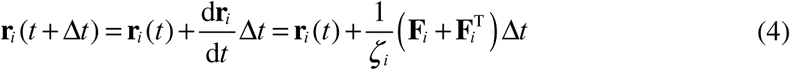

For the deterministic force, we consider i) extensional forces which maintain the equilibrium lengths of segments, ii) bending forces that maintain equilibrium angles formed by segments, iii) torsion forces which maintain equilibrium torsional angles, and iv) repulsive forces which account for volume-exclusion effects between neighboring actin segments. The extensional, bending, and torsional forces are defined by the following harmonic potentials:

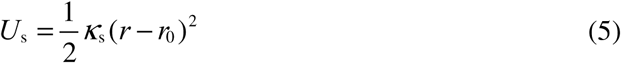

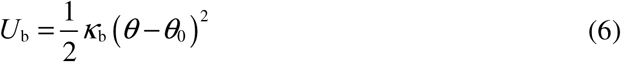

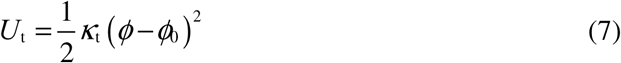

where *κ*_s_, *κ*_b_, and *κ*_t_ represent extensional, bending, and torsional stiffness, respectively. *r* and *r*_0_ are the instantaneous and equilibrium lengths of segments, *θ* and *θ*_0_ are instantaneous and equilibrium angles between interconnected segments, and *φ* and *φ_0_* are instantaneous and equilibrium torsional angles. The repulsive forces acting between neighboring actin filaments are represented by the following harmonic potential (Kim et al., 2009):

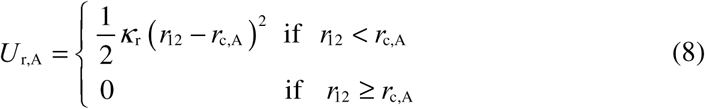

where *κ*_r_ represents the strength of repulsive forces, *r*_12_ is a minimum distance between two neighboring actin segments, and *r*_c,A_ is the diameter of actin segments.

#### Mechanical and geometrical properties of cytoskeletal components

The extensional (*κ*_s,A_) and bending (*κ*_b,A_) stiffnesses of actin segments maintain their equilibrium length (*r*_0,A_ = 140 nm) and equilibrium angle (θ_0,A_ = 0 rad), respectively. Arp2/3 has one extensional stiffness, four bending stiffnesses, and one torsional stiffness. The equilibrium length of two Arp2/3 segments (*r*_0,Arp2/3_ = 35 nm) is maintained by extensional stiffness (*κ*_s,Arp2/3_). An equilibrium angle between two Arp2/3 segments (θ_0,Arp2/3_ = 0 rad) is maintained by bending stiffness (*κ*_b,Arp2/3c_). Two bending stiffnesses (*κ*_b,Arp2/3m_ and *κ*_b,Arp2/3d_) maintain an equilibrium angle between one segment of Arp2/3 and the mother filament (θ_0,Arp2/3m_ = 90° = 1.57 rad) and an equilibrium angle between the other segment of Arp2/3 and the daughter filament (θ_0,Arp2/3d_ = 20° = 0.35 rad), respectively. Another bending stiffness (*κ*_b,Arp2/3f_) maintains an equilibrium angle between the mother and daughter filaments (θ_0,Arp2/3d_ = 70° = 1.22 rad). Torsional stiffness (*κ*_t,Arp2/3_) maintains a zero torsional angle between the mother and daughter filaments to enforce them to exist on a single plane.

### Dynamic behaviors of cytoskeletal components

#### Binding between profilin and actin monomers

Actin monomers in this model can exist in two states: profilin-free actin and profilin-bound actin. The actin monomers undergo a transition between the two states at the rate of *k*_+,prof_ (profilin-free actin to profilin-bound actin) and *k*_-,prof_ (profilin-bound actin to profilin-free actin). The quantitative immunoblotting of intracellular proteins in *Arabidopsis* showed an excess of profilin relative to actin (Chaudhry et al. 2007; Table 1). Thus, we set *k*_+,prof_ significantly higher than *k*_-,prof_ to ensure that >90% of actin monomers exist in the profilin-bound state.

#### De novo nucleation, polymerization, and depolymerization

Actin filaments can be formed by de novo nucleation which is represented by the emergence of one actin segment. The rate of de novo nucleation is determined by a rate constant, *k*_+,n_, and the concentration of profilin-free actin monomers (Blanchoin et al., 2014; Goode et al., 2023; Pollard & Borisy, 2003). Due to very low *k*_n,A_, the de novo nucleation rarely occurs. To reach a steady state faster, 10 filament seeds, each of which corresponds to an actin segment formed by the de novo nucleation, are randomly placed within the domain at the beginning of the simulation.

Then, actin polymerization occurs only from the barbed end of actin filaments with a rate constant, *k*_p,A_, by adding one actin segment. For polymerization, we assume that both profilin-free and profilin-bound actin can be used unlike de novo nucleation. Right after nucleation and polymerization, actin is assumed to be in the ATP state. After the time duration of *t*_ATP_, actin transitions from the ATP state to the ADP state. Actin depolymerization takes place from the pointed end of actin filaments in the ADP state at a constant rate, *k*_d,A_, by removing one actin segment. Both polymerization and depolymerization occur without dependence on forces.

The plant cortical actin array maintains a unique steady state in which the actin polymerization rate is ∼10 times higher than the depolymerization rate (Staiger et al. 2009; S. Zhang et al. 2016). Although such asymmetry between polymerization and depolymerization rates is expected to result in the continuous filament growth, actin filaments within the cortical actin array of plant cells have an average length of 10–17 µm (Table 1). This suggests that severing and capping proteins likely play crucial roles in promoting filament disassembly, helping mitigate the effects of this asymmetry.

#### Severing

To account for the severing activity of ADF/cofilin (Maciver & Hussey, 2002; McCullagh et al., 2014; McGough et al., 1997), actin filaments can be severed at any point in the ADP state at a rate proportional to a local bending angle, θ:

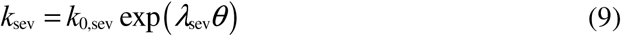

where *k*_0,sev_ and λ_sev_ represent the zero-angle severing rate constant and angle sensitivity, respectively. The severing event corresponds to the disappearance of one actin segment within the filament. We assume that *k*_0,sev_ is extremely low, but λ_sev_is quite high. Thus, a change in θ induced by the thermal fluctuation of actin filaments is not large enough to significantly enhance the severing rate. However, if there are cross-linkers and molecular motors, *k*_sev_ can be much higher than *k*_0,sev_ due to locally large θ. For example, myosin XI is known to cause filament buckling (Cai et al., 2014), which can result in a substantial increase in θ (Duan & Tominaga, 2018). For the purpose of our simulation, we vary only *k*_0,sev_.

#### Capping

Filament capping is modeled as a transition between two states at the barbed end of actin filaments. We assume that formin-bound barbed ends cannot be capped. Immediately after polymerization, the barbed end remains available for polymerization. However, it can transition to a capped state at the rate of *k*_+,cap_, after which polymerization ceases. The reverse transition occurs at the rate of *k*_-,cap_. In this model, *k*_+,cap_ is kept constant, assuming an abundant supply of capping proteins, thereby neglecting their depletion. A recent study suggests that ADF/cofilin not only promotes severing but also restricts polymerization at barbed ends newly formed after severing (Michelot et al. 2006). Based on this finding, we assume that barbed ends formed after severing events are immediately capped, enhancing F-actin disassembly.

#### Branch formation by Arp2/3 complex

In this model, we consider a side-branching nucleator as an explicit element, to represent Arp2/3 complex. It is assumed that Arp2/3 can bind to any part of a mother filament with the rate constant of *k*_+,Arp2/3_. Immediately after the binding event, one actin segment (either of profilin-free actin or profilin-bound actin) is added to the other side of Arp2/3 to mimic Arp2/3-induced branch nucleation. After the nucleation, the branch is elongated by actin polymerization, similar to other filaments formed by de novo nucleation. Considering that the dissociation of Arp2/3 from the daughter filament occurs more readily than from the mother filament (L. Y. Cao & Way, 2024; Ghasemi et al., 2024), we assign distinct unbinding rates: *k*_-,Arp2/3m_ for the mother and *k*_-,Arp2/3d_ for the daughter filament.

#### Enhanced polymerization by processive formin

It is assumed that formin binds to the free (i.e., uncapped) barbed end of actin filaments to increase the polymerization rate 3 times, based on the experimental finding that reported ∼2–3 fold enhanced elongation rate measured at the formin-bound barbed end (S. Zhang et al. 2016; Table 1). For implementing the formin activity, actin located at the barbed end in the model can undergo a transition between two states: a free state and a formin-bound state. The transition from the free state to the formin-bound state occurs at the rate of *k*_+,form_. After the transition, the barbed end returns to the free state after the time duration of *t*_form_. We assume that a barbed end in the formin-bound state cannot simultaneously transition to a capped state.

### Measurement, analysis, and visualization

The number of actin in each of 5 states is counted once per second: i) profilin-bound monomer, ii) profilin-free monomer, iii) formin-elongated filament, iv) Arp2/3-formed filament, and v) filament formed by de novo nucleation. Note that the formin-elongated filament may originate from either Arp2/3-mediated nucleation or de novo nucleation. The number and average length (in μm) of filaments formed by Arp2/3 or de novo nucleation are quantified once per second. Polymerization rates are calculated in μm/s by measuring an increase in average filament length with a 100-second interval. Only free (uncapped) barbed ends are considered for this calculation. Depolymerization rates are calculated (in μm) by measuring a decrease in average filament length with a 100-second interval. Severing rates are calculated in events/μm/s by dividing the number of severing events with a 100-second interval by the average filament length and the time interval. The number of capped barbed ends is counted once per second.

Networks are visualized using Visual Molecular Dynamics (VMD, University of Illinois at Urbana-Champaign). Actin filaments are marked using different colors, depending on whether it is formed by de novo nucleation, Arp2/3-mediated nucleation, and formin-enhanced polymerization. Arp2/3 is represented by small green spheres.

### Computational setup

To simulate the plant cortical actin array, we employ a three-dimensional computational domain (20×20×1 µm). The thickness of the domain in the z direction is assumed to be small due to the central vacuole within the plant cytosol, which can occupy up to 90% of the cellular volume (Festa et al., 2016). Periodic boundary conditions are applied only in the x and y-directions, whereas two boundaries normal to the z-direction exert repulsive forces to elements that are displaced out of the domain (Fig. 2). Network assembly is mediated by the de novo nucleation, spontaneous polymerization, formin-induced enhanced polymerization, and branching of actin filaments, and network disassembly is induced by the depolymerization, severing, and capping of actin filaments. We performed simulations for 2000 s with a variation in key parameters relative to the reference condition, including *k*_0,sev_, *k*_+,cap_, *k*_+,form_, *k*_+,prof_, and the molar ratio of Arp2/3 (*R*_Arp2/3_). Under the reference condition, total actin concentration is 1 µM, and *R*_Arp2/3_ is 0.00007. The reference values of all other parameters are listed in Table 2.

## AUTHOR CONTRIBUTIONS

J.H.K., C.J.S, and T.K. designed the project. J.H.K. performed simulations and analyzed data obtained from the simulations. T.K. supervised all computational studies. W.Z. prepared experimental images, and C.J.S. supervised all experimental studies. All of the authors participated in discussing the data and writing the manuscript.

## ACKNOWLEDGMENTS

We gratefully acknowledge the support from the EMBRIO Institute, contract #2120200, a National Science Foundation (NSF) Biology Integration Institute.

## CONFLICT OF INTEREST DISCOSURE

The authors declare no conflict of interest.

## DATA AVAILABILITY STATEMENT

The data that support the findings of this study are available from the corresponding authors upon reasonable request.

